# Catalase protects against non-enzymatic decarboxylations during photorespiration

**DOI:** 10.1101/2021.08.04.455084

**Authors:** Han Bao, Matt Morency, Winda Rianti, Sompop Saeheng, Sanja Roje, Andreas P.M. Weber, Berkley Walker

## Abstract

Photorespiration recovers carbon that would be otherwise lost following the oxygenation reaction of rubisco and production of glycolate. Photorespiration is essential in plants and recycles glycolate into usable metabolic products through reactions spanning the chloroplast, mitochondrion, and peroxisome. Catalase in peroxisomes plays an important role in this process by disproportionating H_2_O_2_ resulting from glycolate oxidation into O_2_ and water. We hypothesize that catalase in the peroxisome also protects against non-enzymatic decarboxylations between hydrogen peroxide and photorespiratory intermediates (glyoxylate and/or hydroxypyruvate). We test this hypothesis by detailed gas exchange and biochemical analysis of *Arabidopsis thaliana* mutants lacking peroxisomal catalase. Our results strongly support this hypothesis, with catalase mutants showing gas exchange evidence for an increased stoichiometry of CO_2_ release from photorespiration, specifically an increase in the CO_2_ compensation point, a photorespiratory-dependent decrease in the quantum efficiency of CO_2_ assimilation, increase in the ^12^CO_2_ released in a ^13^CO_2_ background and an increase in the post-illumination CO_2_ burst. Further metabolic evidence suggests this excess CO_2_ release occurred via the non-enzymatic decarboxylation of hydroxypyruvate. Specifically, the catalase mutant showed an accumulation of photorespiratory intermediates during a transient increase in rubisco oxygenation consistent with this hypothesis. Additionally, end products of alternative hypotheses explaining this excess release were similar between wild type and catalase mutants. Furthermore, the calculated rate of hydroxypyruvate decarboxylation in catalase mutant is much higher than that of glyoxylate decarboxylation. This work provides evidence that these non-enzymatic decarboxylation reactions, predominately hydroxypyruvate decarboxylation, can occur *in vivo* when photorespiratory metabolism is genetically disrupted.

**One Sentence Summary:** Catalase guards against additional carbon loss from photorespiration arising from non-enzymatic decarboxylations of photorespiratory intermediates.

## Introduction

Photorespiration is the single largest limitation to C3 photosynthesis under current atmospheres, consuming ∼30-40% of total plant energy in the light and resulting in rates of CO_2_ loss approaching 25% the rate of net CO_2_ fixation (Sharkey 1988; Walker et al. 2016b). Given this major role in determining net rates of energy use and CO_2_ exchange, it is vital to understand the biochemical underpinnings of photorespiration to both accurately model plant productivity in response to changing climates and design optimization strategies for improving net photosynthesis. Improvement strategies targeting photorespiration have been showing initial promise both under laboratory conditions (Timm et al. 2012) and more recently under in-field experiments (South et al. 2019), however; future efforts in optimization and improved modeling may require a more mechanistic understanding of the function of native photorespiration.

Photorespiration recycles 2-phosphoglycolate (2-PG) produced following the reaction of ribulose 1,5-bisphosphate (RuBP) with O_2_ as catalyzed by the first enzyme of the C3 cycle, RuBP carboxylase/oxygenase (rubisco). This recycling pathway comprises over a dozen enzymatic conversions and transport steps spanning the chloroplast, peroxisome and mitochondria and results in the partial recycling of 2-PG into the C3-cycle intermediate 3-phosphoglycerate (3-PGA) with the loss of CO_2_ and energy (Figure 1, (Foyer and Noctor 2009; Bauwe et al. 2010). The CO_2_ loss from photorespiration is assumed to come primarily from glycine decarboxylation in the mitochondria, resulting in a stoichiometric release of 0.5 CO_2_ per rubisco oxygenation (Somerville and Ogren 1980; Somerville 2001; Abadie et al. 2016). The stoichiometric release of CO_2_ per rubisco oxygenation is a cornerstone assumption for biochemical models of leaf photosynthesis, which represent net CO_2_ fixation rates in scales ranging from the single cell to the entire globe (Farquhar et al. 1980; von Caemmerer and Farquhar 1981; von Caemmerer 2013; Sun et al. 2014).

**Figure 1.**
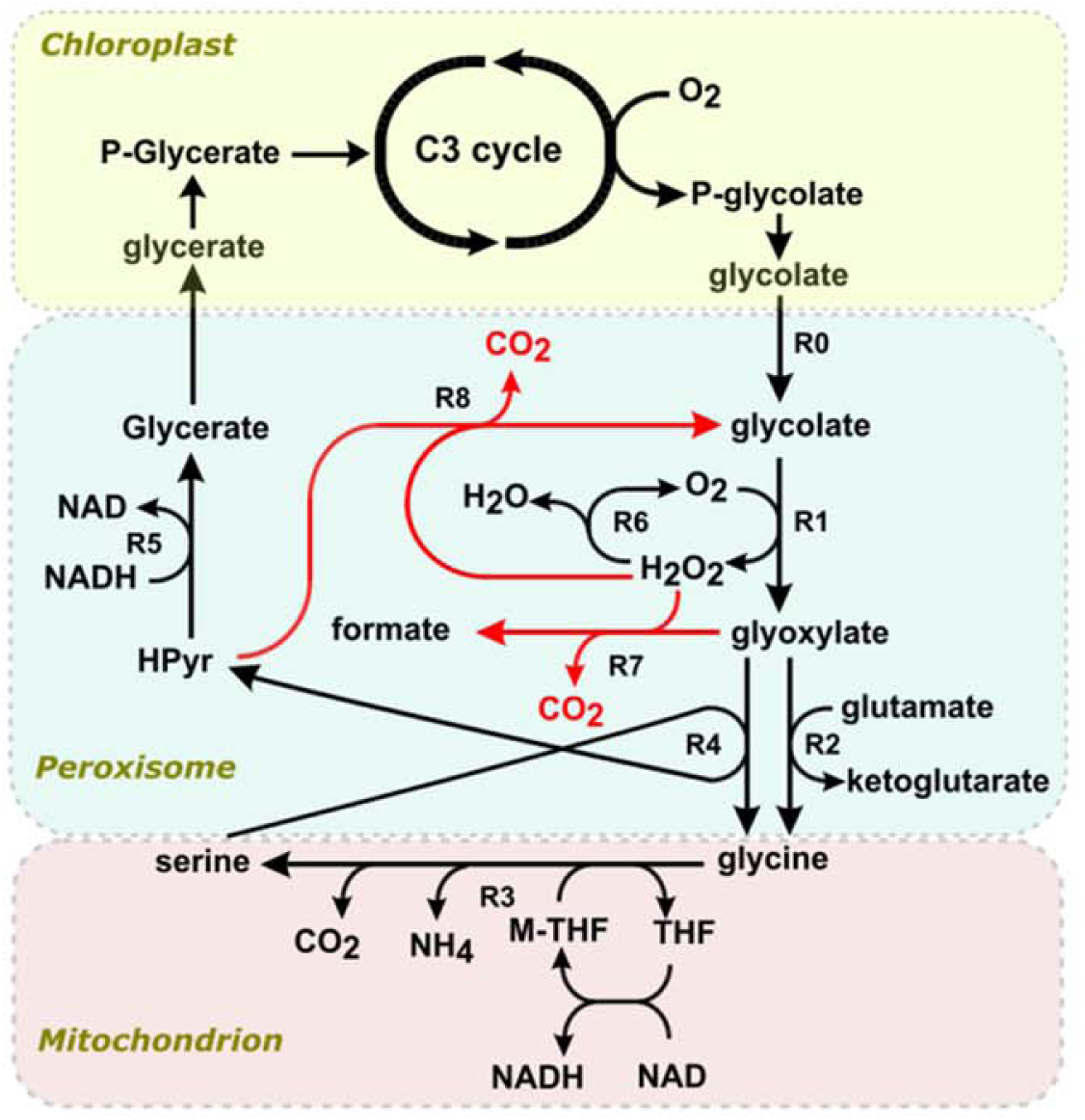
Schematic representation of photorespiration model developed in this work. Enzymatic reactions (R1-R6) typically associated with photorespiration are shown in black while non-enzymatic decarboxylation reactions (R7-R8) are shown in red. RO represents transport of glycolate into the peroxisomes. Glycolate oxidase (Rl) catalyzes the conversion of glycolate to glyoxylate and H_2_O_2_. The latter molecule is decomposed to oxygen and water by catalase (R6). Glyoxylate is aminated by either glutamate glyoxylate aminotransferase (R2) or serine glyoxylate transaminase (R4) to produce glycine. Glycine is then decarboxylated to produce serine while releasing CO_2_, NH_4_, cycling tetrahydrofolate (THF) and methyl-THF (M-THF) and reducing NAD by a multienzyme complex, glycine-cleavage system, in the mitochondria. This conversion of glycine to serine is modelled as a single reaction (R3) where flux in R3 equally contributes to serine and CO_2_ formation. Hydroxypyruvate (HPyr) produced through serine glyoxylate transaminase (R4) is reduced by hydroxypyruvate reductase (R5) to form glycerate, which is transported back to the chloroplast for incorporation into the C3 cycle. Non-enzymatic decarboxylations can occur either between glyoxylate and H_2_O_2_ (R7) or between HPyr and H_2_O_2_ (R8), both releasing CO_2_ in the process. Note that the process of photorespiration involves different compartments however, compartmentalization has not been taken into consideration in this model. The concentrations of the intermediates NAD, NADH, glutamate, THF, M-THF and O_2_ are assumed to be present in excess to drive the associated reaction at maximal rate.

While there is strong evidence that glycine decarboxylation is the predominate source of CO_2_ loss from photorespiration, there are other potential reactions that can result in additional CO_2_ loss including the non-enzymatic decarboxylation (NED) of glyoxylate (Halliwell and Butt 1974; Grodzinski 1978) and/or hydroxypyruvate by H_2_O_2_ (Cousins et al. 2008; Keech et al. 2012). NED reactions would reduce the carbon recycling efficiency of photorespiration by increasing the stoichiometric release of CO_2_ per rubisco oxygenation by up to 400%, assuming they processed all of the photorespiratory flux (Cousins et al. 2011). While early *in vitro* experiments offered support for the importance of NED reactions in explaining *in vivo* CO_2_ loss in wild type (WT) plants (Halliwell and Butt 1974; Grodzinski and Butt 1976; Grodzinski 1978), subsequent genetic and flux labeling experiments demonstrate that glycine decarboxylation explains the majority of CO_2_ loss *in vivo*, at least under ambient (20-25 °C) conditions (Somerville and Ogren 1980; Somerville 2001; Abadie et al. 2016). These findings indicate that catalase activity, which detoxifies H_2_O_2_ in the peroxisome, may be present in high enough levels to inhibit NED reactions under the conditions measured in WT plants, at least under ambient temperatures.

When photorespiration is disrupted genetically, however, there is evidence that NED reactions drive excess carbon loss. For example, *hpr* mutants lacking peroxisomal hydroxypyruvate reductase (HPR) have increased photorespiratory CO_2_ compensation points (Γ*) and release of CO_2_, (Cousins et al. 2008; Cousins et al. 2011; Timm et al. 2008; Keech et al. 2012), demonstrating an increase in the stoichiometry of CO_2_ released per rubisco oxygenation (See Theory). Additionally, mutants lacking the foliar-expressed catalase (CAT) isoform (*cat2*) similarly showed increases in the compensation point (Γ) and other gas exchange signatures of CO_2_ release from photorespiration, but this was measured under a limited set of conditions and not assessed using approaches that allow measurements under ambient CO_2_ concentrations (Keech et al. 2012). Furthermore, additional evidence is needed to establish *which* NED reactions (glyoxylate or hydroxypyruvate decarboxylation) explain this *in vivo* instance of excess carbon loss from photorespiration and that catalase specifically plays a critical role in protecting against this excess loss in WT plants.

In this work, we present gas exchange and metabolic data to demonstrate that catalase mutants show an increase in the stoichiometry of CO_2_ release per rubisco oxygenation and this excess CO_2_ release from photorespiration most likely comes from decarboxylation of hydroxypyruvate by H_2_O_2_. These findings suggest catalase plays a critical role in guarding against additional wasteful loss of CO_2_ from photorespiration and provide a set of approaches that could be used to examine the mechanisms governing the efficiency of CO_2_ release from photorespiration under elevated temperatures in WT plants.

## Theory

### Gas exchange theory

The presence of additional decarboxylation reactions during photorespiration was examined using various gas exchange approaches; specifically, measurements of Γ*, Γ, and the quantum efficiency of CO_2_ fixation (Φ_CO2_) measured under differing photorespiratory conditions. Each of these measurements are impacted by the amount of CO_2_ released per rubisco oxygenation from photorespiration *in vivo* as described below.

Γ* is a key parameter that links plant biochemistry to rates of net gas exchange by combining rubisco specificity for reaction of CO_2_ relative to O_2_ (S_c/o_) with oxygen concentration (O) and the amount of CO_2_ lost from photorespiration per rubisco oxygenation (η) according to

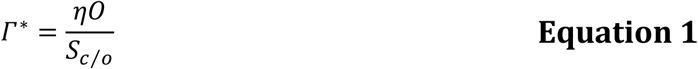

(Farquhar et al. 1980; von Caemmerer and Farquhar 1981; von Caemmerer 2013). Since Γ* is proportional to η and NED reactions increase η, differences in Γ* can indicate changes in the amount of CO_2_ released from photorespiration due to increases in NED reactions, especially when measurements are done under the same O and in the same species with identical S_c/o_.

Γ is a more readily measured parameter, but is more indirectly related to η since it measures the CO_2_ compensation point where a leaf assimilates as much CO_2_ as it releases from both photorespiration and non-photorespiratory CO_2_ loss in the light (R_L_) according to

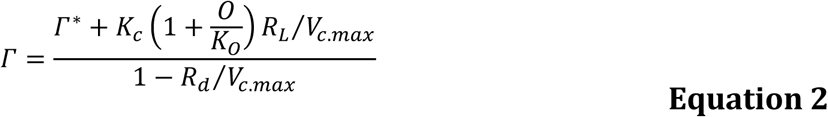

where *K*_c_, *K*_o_, *R*_L_ and *V*_c,max_ are the Michaelis-Menten enzyme constants of Rubisco for CO_2_, O_2_, rate of mitochondrial respiration in the day and maximum rate of Rubisco carboxylation. Since Γ is also dependent on Γ*, it is similarly impacted by changes in η driven by NED reactions.

A third test for the presence of NED reactions during photorespiration is in measurements of Φ_CO2_. Since Φ_CO2_ represents net CO_2_ fixation per absorbed photon of light, Φ_CO2_ should decrease when total amounts of CO_2_ lost from photorespiration increase and reduce net CO_2_ fixation. The total rate of CO_2_ lost from photorespiration is equal to η multiplied by rates of rubisco oxygenation (*V*_*o*_), so *V*_*o*_ dependent decreases in Φ_CO2_ would provide further evidence for the presence of NED reactions.

## Results

### Measurements of Γ* and Γ were higher in *cat2*

To determine if *cat2* had elevated compensation points consistent with an increase in CO_2_ release per rubisco oxygenation, Γ* and Γ were measured using the common intersection method (Walker and Ort 2015; Walker et al. 2016a). Γ^*^ in *cat2* is 30% greater than in the WT under 25 °C (Supplemental Figure 1). This increase in Γ^*^ corresponds to an increase in CO_2_ release per rubisco oxygenation from 0.5 to 0.64, assuming S_c/o_ stays constant (Equation 1). Furthermore, Γ was significantly higher for every light intensity used during the common intercept measurement of Γ* except under the lowest light intensity (50 μmol m^-2^ s^-1^), suggesting that the impact of deficient CAT activity is greatest under elevated rates of photorespiration (Supplemental Table I).

### *cat2* had a lower efficiency of net carbon assimilation under higher rates of photorespiration

The response of net assimilation to light intensity was used to determine if *cat2* had a photorespiratory-dependent decrease in Φ_CO2_ driven by increases in CO_2_ release per rubisco oxygenation. Consistent with this hypothesis, Φ_CO2_ was much lower in *cat2* compared to the WT plants under the highest rates of rubisco oxygenation (*v*_*o*_), but not when *v*_*o*_ was low as measured under high CO_2_ or low O_2_ (Figure 2). Furthermore, the decrease in Φ_CO2_ of *cat2* compared to the WT plants followed a roughly linear trend with *v*_*o*_. This trend is consistent when Φ_CO2_ is compared to the ratio *v*_*o*_/*v*_*c*_, with higher ratios in *cat2* showing lower efficiencies. Interestingly, under very high rates of photorespiration, *cat2* actually *loses* more CO_2_ than is fixed as light intensity increases, resulting in a negative Φ_CO2_ (under 5 Pa CO_2_, Figure 2 and Supplemental Figure 2).

**Figure 2.**
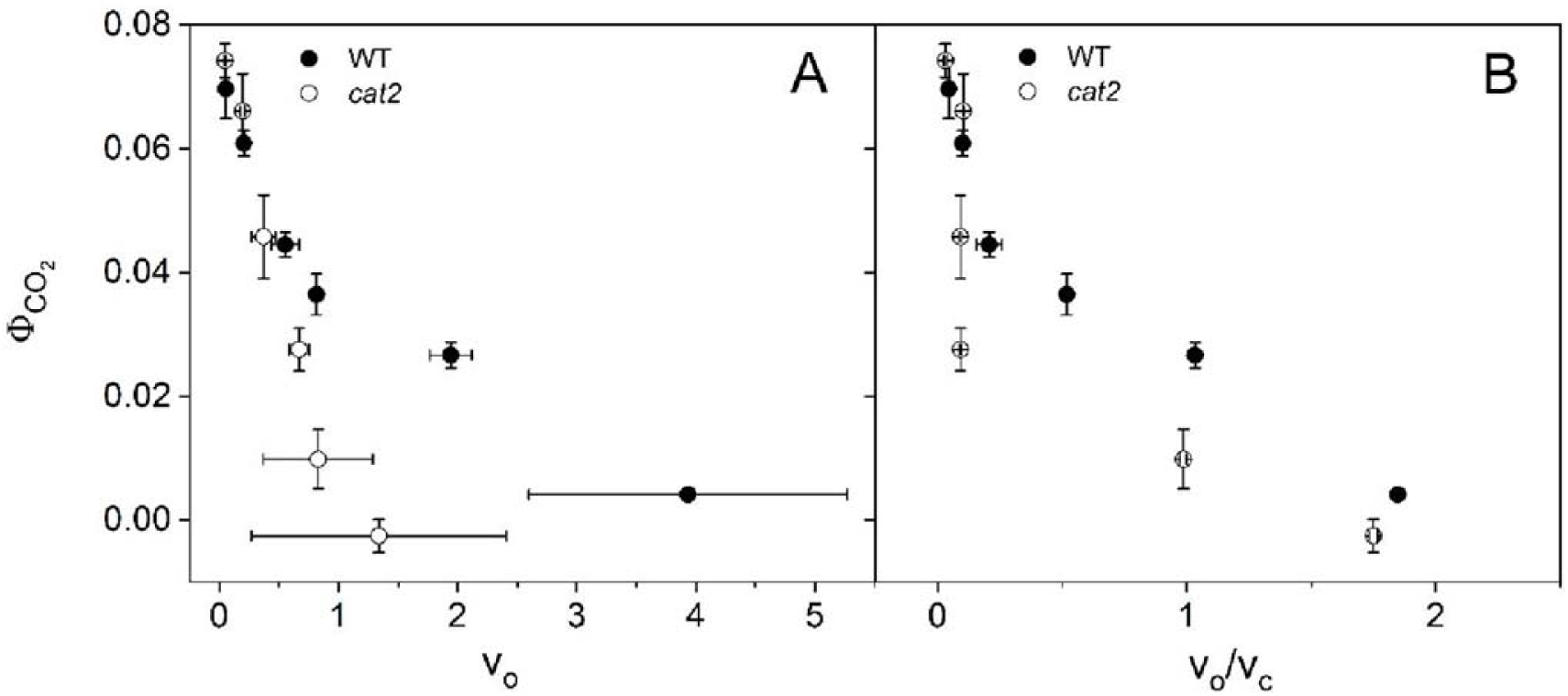
The response of the quantum efficiency of CO_2_ fixation (Ф_CO2_) to different rates of Rubisco oxygenation (v_o_, A) and the ratio of rubisco oxygenation to carboxylation (v_o_/ v_c_, B) in *A. thaliana* wild type (WT) and plants lacking peroxisomal-type catalase expression *(cat2)*. Ф_CO2_ was determined from the initial slopes of light response curves under various CO_2_ and O_2_ partial pressures measured using the LI-COR 6800 infra-red gas analyzer. Shown with n=5 ± ster.

Fluorescence measurements indicate that decreases in Φ_CO2_ were not due to general damage or inhibition to the core photosynthetic machinery of *cat2*, but indeed photorespiratory-dependent. For example, *cat2* and WT have similar dark and light adapted values of the quantum yield of photosystem II (F_v_/F_m_ and F_v_’/F_m_’) and rates of non-photochemical quenching (NPQ, Supplemental Table II).

To determine if other, non-photorespiratory rates of CO_2_ release changed with photorespiratory conditions to reduce Φ_CO2_ we compared measurements of R_L_ between WT and *cat2* under various CO_2_ concentrations and O_2_ concentrations. In all cases, there was either no significant difference or there was a slightly lower R_L_ in *cat2*, suggesting that the observed decreases in Φ_CO2_ are unlikely due to changes in non-photorespiratory CO_2_ release (Supplementary Figure 3).

### The stoichiometry of CO_2_ released per rubisco oxygenation increases in *cat2*

To confirm more directly that the *cat2* plants had an increase in CO_2_ release per rubisco oxygenation under higher light intensities, we next used membrane-inlet mass spectroscopy to determine relative rates of CO_2_ release from photorespiration per rubisco oxygenation. This was necessary since Γ* and Φ_CO2_ are both measured under low light intensities due to the nature of the gas exchange approaches and we wanted to confirm this evidence for increased CO_2_ release under more physiological conditions. Membrane-inlet mass spectroscopy determines rates of rubisco oxygenation and relative rates of CO_2_ release from photorespiration from net fluxes of O_2_ and CO_2_ resolved from photosynthesis using isotopically enriched atmospheres (Cousins et al. 2008; Canvin et al. 1980). Measurements of the total ^12^CO_2_ released in the light under a saturating concentration of ^13^CO_2_ were higher in *cat2*, indicating that *cat2* had a higher efflux of CO_2_ compared to WT (Supplemental Figure 4a). To ensure this release was not due simply to higher rates of rubisco oxygenation, and a true increase in the stoichiometric release of CO_2_ per oxygenation, ^13^CO_2_ release was normalized by rates of rubisco oxygenation (Supplemental Figure 4b). This normalization also showed that the stoichiometric release of CO_2_ per rubisco oxygenation was higher in *cat2* compared to WT, consistent with additional non-enzymatic decarboxylation reactions from photorespiration.

### Evidence for higher and alternate sources of CO_2_ release from the post-illumination burst and photorespiratory isotopic fractionation

To determine how prevalent these increases in CO_2_ release in *cat2* are under ambient CO_2_ concentrations, we next measured the post-illumination burst (PIB). The PIB refers to the burst of CO_2_ released from leaves immediately after a light to dark transition (Bulley and Tregunna 1971; Doehlert et al. 1979). Although the PIB is not a strictly quantitative measurement, it can be used to estimate the amount of CO_2_ release from photorespiration. Our measurements of the PIB revealed that the total CO_2_ release following a period of illumination was higher in *cat2* as compared to WT (Figure 3a). Furthermore, the PIB peak was integrated to determine the magnitude of total CO_2_ release during PIB. The data shows that the CO_2_ evolution in *cat2* was nearly two-fold greater than WT (Figure 3b), reflecting an increased stoichiometry of CO_2_ release per rubisco oxygenation.

**Figure 3.**
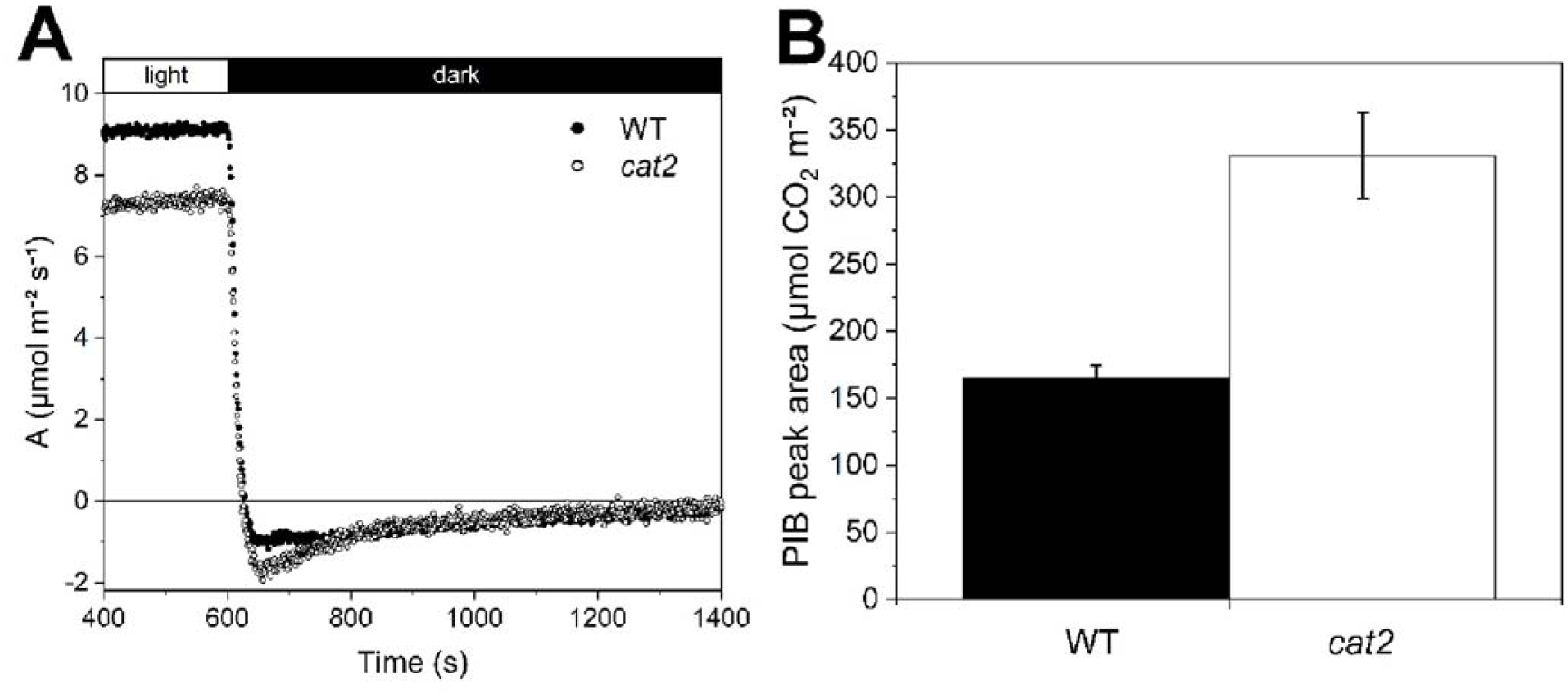
Post-Illumination Burst of CO_2_ (PIB) in *A. thaliana* wild type (WT) and plants lacking peroxisomal-type catalase expression *(cat2)*. (A) Changes in the rate of uptake and release of CO_2_ were measured during a 40-min light period followed by a 20-min dark period using the Ll-COR 6800 infra-red gas analyzer. (B) Quantification of PIB by integration of peak area of CO_2_ release. Shown with n=5 ± ster.

### Metabolic transients, formate and folic acid concentrations suggest hydroxypyruvate decarboxylation releases excess CO_2_ in *cat2*

To resolve the origin of this excess CO_2_ release from photorespiration, we next examined the response of photorespiratory metabolites during a transient period of increasing photorespiration induced by measuring a plant switching from a low to high light condition. Metabolite concentrations can be more informative when measured under transient conditions, before a new steady-state is established (Abadie et al. 2016). Specifically, we hypothesized that if glyoxylate NED explained this excess release, more carbon would leave photorespiration in the form of formate, decreasing the pool sizes of intermediates downstream of glyoxylate (glycine, serine, hydroxypyruvate and glycerate) during transient increases in photorespiratory flux in *cat2* as compared to WT (Figure 1). Alternatively, hydroxypyruvate NED forms the photorespiratory intermediate glycolate, meaning that this NED reaction should result in relative increases in photorespiratory intermediate pools (for a given change in v_o_) as carbon is maintained in the cycle in *cat2* as compared to WT.

Our transient time-course measurements showed higher relative pool sizes of photorespiratory intermediates in *cat2*, supporting NED of hydroxypyruvate as the source of excess CO_2_ release (Figure 4). Specifically, all photorespiratory metabolites increased more with increased photorespiratory rates in *cat2*, except for glyoxylate (Figure 4). This general trend is consistent with more carbon staying within the photorespiratory cycle in *cat2*, as expected with the NED of hydroxypyruvate to glycolate, which would maintain more carbon in photorespiration as opposed to converting hydroxypyruvate to glycerate (Figure 1). These general trends, however, could be explained if *cat2* had higher relative rates of v_o_ during this light transient.

**Figure 4.**
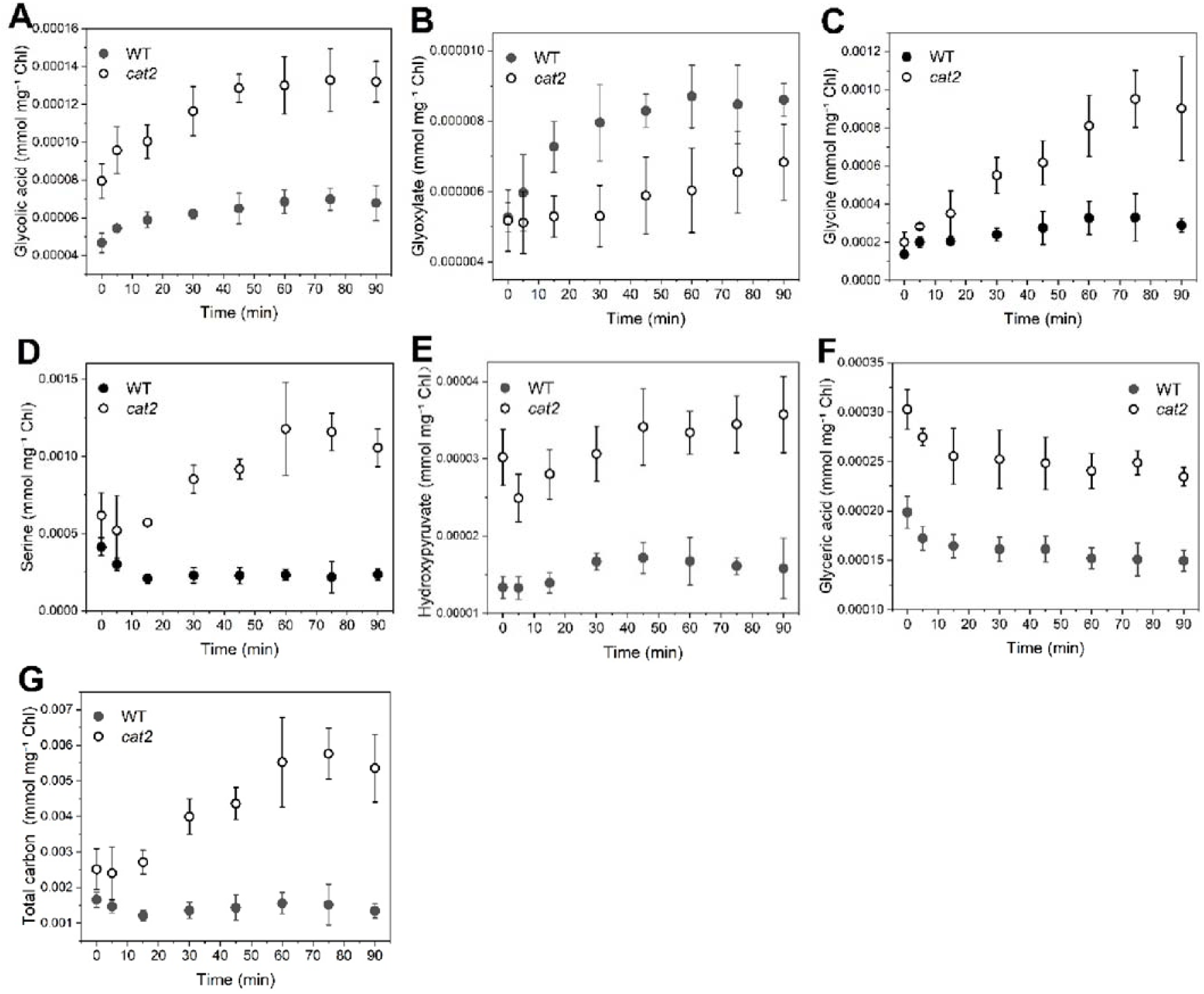
Metabolic changes in photorespiration upon transfer from low to high light for *A. thaliona* wild type (WT) and plants lacking peroxisomal-type catalase expression *(cat2)*. Irradiance changes were from 50 to 400 μE m^-2^ s^-1^. Concentrations of glycolate (A), glyoxylate (B), glycine (C), serine (D), Hydroxypyruvate (E) and glycerate (F) were detected by GC-MS. Total carbon concentrations (G) were calculated based on the above concentrations except for glycerate. The data are given as mean (n=5) ± ster.

Complementary gas exchange data showed that these differences in metabolite responses are not associated with a higher rate of v_o_ in *cat2*. The rates of v_o_ increased immediately upon exposure to high light, as shown by v_o_ estimated from gas exchange measurements made over a similar period (Supplemental Figure 5). Moreover, the two light induction curves were almost identical, indicating similar rates of glycolate influx between WT and *cat2*. However, WT and *cat2* had very different metabolic responses during the photorespiratory transient. For example, the pool size of glycolate in c*at2* had a greater proportional increase than that in WT (Figure 4A). Similar trends were also observed for glycine, serine and hydroxypyruvate (Figure 4C, D, E), while the opposite trend was seen for glyoxylate (Figure 4B).

The above results show how each individual metabolite responded to the transient but to understand how much total carbon was present in the photorespiratory intermediates at each timepoint, we determined the total carbon within photorespiration by summing the carbon present in each individual metabolite. We calculated total carbon concentration based on the five metabolite pools between glycolate and hydroxypyruvate (Figure 4G, glycolate, glyoxylate, glycine, serine, and hydroxypyruvate). These carbon pools should increase in total in the presence of hydroxypyruvate NED during an increase in photorespiration. Our data showed that, compared to a relatively flat response curve in WT, a larger amount of carbon accumulated in *cat2* as compared to WT during the transient period, consistent with this hypothesis (Figure 4G).

Formate is a product of glyoxylate NED. Formate can either be decarboxylated in the mitochondria, or enter one carbon metabolism (a cycle involving numerous folate species) following a reaction catalyzed by tetrahydrofolate ligase (Hanson and Roje 2001). To further test if this NED reaction also contributes to the excess carbon loss, we measured the contents of formate and its downstream folate species. If glyoxylate decarboxylation takes place *in vivo*, we might expect to see a higher level of formate and/or folates in *cat2*. However, our data shows that there is no significant difference between WT and *cat2* in formate or folate contents (Supplemental Figure 6), suggesting that the glyoxylate decarboxylation is not the predominant NED reaction under physiological conditions.

### Determining H_2_O_2_ concentrations

To determine if the metabolite concentrations that we measured were large enough to drive high rates of particular NED’s, we needed to measure levels of H_2_O_2_ in WT and *cat2* during the transient from low to high photorespiratory rates explored above. There is no difference in H_2_O_2_ concentrations measured under the initially low photorespiratory rates (time t=0), implying that *cat2* has adapted to the stress of decreased catalase by activating alternate antioxidative systems for H_2_O_2_ scavenging to compensate for the shortage in catalase. However, a large divergence was observed after a shift to high light. The H_2_O_2_ level was elevated by ∼50% in *cat2* and was reduced by ∼40% in WT by the end of the transient (Figure 5a). We hypothesize that the decrease in H_2_O_2_ concentration under elevated light in WT might be due to the light activation of the catalase enzyme, but regardless, *cat2* plants had elevated H_2_O_2_ content, a key substrate for NED’s. To determine if the content of H_2_O_2_ was large enough to drive NED, we next needed some additional parameterizations of the reaction.

**Figure 5.**
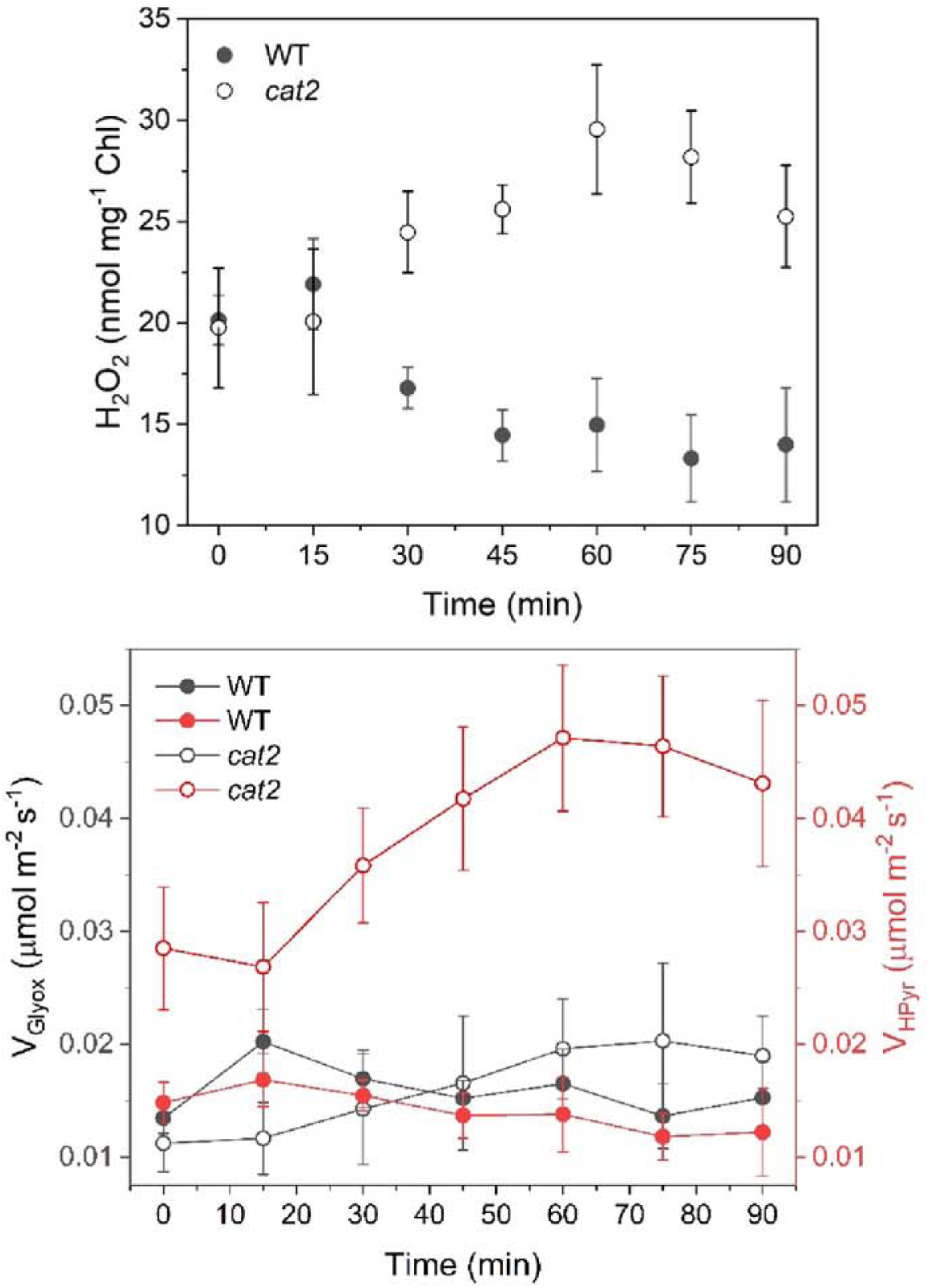
Hydrogen peroxide (H_2_O_2_) changes in *A. thaliana* wild type (WT) and plants lacking peroxisomal-type catalase expression *(cat2)* upon transfer from low to high light (A) and comparison of reaction rates of glyoxylate decarboxylation (black) and hydroxypyruvate decarboxylation (red) in *A. thaliana* WT *cat2* (B). Concentrations of substrate (H_2_O_2_, glyoxylate and hydroxypyruvate) during transition from low to high light were the same as shown in Fig. 4 for calculation of decarboxylation rates. Rate constants and order were determined as described in Material and Method section. Data are expressed as mean (n=3-5) ± ster.

### Parameterizing rate constants

To determine if the metabolite concentrations that we measured are large enough to drive high rates of NED’s we characterized the rate constants of the second-order reactions between H_2_O_2_ and either glyoxylate and hydroxypyruvate. To characterize the NED’s reaction order and rate constants, we measured the decay of H_2_O_2_ and substrate following reaction with hydroxypyruvate or glyoxylate using UV spectroscopy at various concentrations of each reactant (Supplemental Figure 7). The response of the reaction rate was linearly related both to [H_2_O_2_] and either [glyoxylate] or [hydroxypyruvate], confirming that both reactions are described by a second-order rate equation. The rate constant for decarboxylation of glyoxylate with H_2_O_2_ (7.5 L mol^-1^ s^-1^) was higher than that describing reaction with hydroxypyruvate and H_2_O_2_ (3.26 mol-1 s^-1^).

### Estimating reaction rates of NED

The reaction rates of NED were determined by multiplying the second order rate constant by the molar concentrations of the reactants (H_2_O_2_ and glyoxylate/hydroxypyruvate). Among the four NED reactions (two in WT and two *cat2*), hydroxypyruvate decarboxylation in *cat2* has the highest rate throughout the transient period (Figure 5b). The rate of hydroxypyruvate decarboxylation in *cat2* was approximately 2 to 5-fold greater than that of the other reactions.

The rates of this CO_2_ loss estimated from hydroxypyruvate NED was ∼5 fold lower than the excess CO_2_ loss predicted from our gas exchange measurements when expressed on a leaf area basis. Specifically, the metabolite data estimates a loss of 0.03 to 0.05 μmol m^-2^ s^-1^, but the gas exchange measurements suggest an excess rate of CO_2_ release of 0.09 to 0.35 μmol m^-2^ s^-1^. We attribute this discrepancy to underestimates of the highly reactive H_2_O_2_ measured in our leaf tissues, a species that displays large variation in absolute values depending on study and assay technique (Queval et al. 2008). These results provide further evidence supporting the hydroxypyruvate decarboxylation as the predominant NED reaction and source of excess CO_2_ release in *cat2*.

### Decarboxylation of Serine

Besides decarboxylation reactions of photorespiration, there is one other photorespiratory-linked decarboxylation reaction that could release CO_2_. Serine decarboxylase catalyzes the conversion of serine to ethanolamine. Phosphorylated ethanolamine is the precursor for the biosynthesis of polar head groups of two phospholipids classes, phosphatidylcholine (PC) and phosphatidylethanolamine (PE). Since *cat2* had a larger pool size of serine (Figure 4D), we wondered whether this could drive a higher rate of serine decarboxylation. To test this hypothesis, we analyzed contents of PC and PE. Our data showed that there were no significant differences in the amount of PC or/and PE between WT and *cat2* (Figure 8A). Furthermore, fatty acid profiles of PC and PE were also similar (Figure 8B). These results further confirm that decarboxylation of hydroxypyruvate is unlikely to be the only predominant source of excess release of CO_2_ in *cat2*.

### Total catalase activity

To confirm and quantify the decrease in catalase activity in *cat2*, total catalase activity was measured via O_2_ evolution. Catalase activity decreased in *cat2* by almost 80% when expressed both on a leaf area and protein content basis (Table I). Furthermore, the decrease in catalase activity was not accompanied by a decrease in total protein content.

**Table I:** Metabolite content, catalase activity and protein content in *A. thaliana* wild type (WT) and plants lacking peroxisomal type catalase expression (*cat2*). Catalase activity was determined from leaf extracts by following the rate of oxygen production following H_2_O_2_ addition in an O_2_ electrode and presented both on a leaf area and mg protein basis. Protein content was determined on the same extract using a Bradford assay of soluble protein. Shown are the averages biological replicates (n=5 ± ster) determined with technical replicates (n=3) with significant differences (students t-test, α < 0.05) indicated by different letters.

|  | WT | *cat2* |
| --- | --- | --- |
| Catalase activity (mol O<sub>2</sub> m<sup>-2</sup> s<sup>-1</sup>) | 0.81±0.09<sup>a</sup> | 0.18±0.02<sup>b</sup> |
| Catalase activity (μmol O<sub>2</sub> mg<sup>-2</sup> Prot s<sup>-1</sup>) | 1.94±0.18<sup>a</sup> | 0.45±0.05<sup>b</sup> |
| Protein content (mg m<sup>-2</sup>) | 415±34<sup>a</sup> | 427±8<sup>a</sup> |

## Discussion

In this paper, we demonstrate that catalase protects against excess photorespiratory carbon loss and that this excess loss decreases net photosynthesis under ambient conditions. In *cat2* plants, both Γ* and Γ were greater than in WT, which is explained by an increase in CO_2_ release per rubisco oxygenation (Supplementary Figure 1 and Supplemental Table I). Additionally, *cat2* Φ_CO2_ had a *V*_*o*_-dependent decrease, which even became negative under high *V*_*o*_ indicating an extra loss of CO_2_ that negatively impacted net photosynthesis and scaled with rates of photorespiration (Figure 2 and Supplemental Figure 1). Furthermore, *cat2* mutants had elevated ^12^CO_2_ release per rubisco oxygenation and a higher PIB peak area than WT, indicating a higher amount of CO_2_ being released under physiological conditions (Supplemental Figure 4 and Figure 3).

Our metabolite data strongly suggest that the source of this excess CO_2_ release from photorespiration arises from the NED reaction between H_2_O_2_ and hydroxypyruvate. Specifically, a larger carbon accumulation of photorespiratory intermediates formed in *cat2* mutant compared to WT during a period of increased photorespiration, suggesting a cyclic route of metabolic flux through photorespiration with NED decarboxylation of hydroxypyruvate to glycolate (Figure 4). Additionally, rates of hydroxypyruvate NED reactions predominated when calculated from measured metabolite pools and reaction rates, providing further evidence for the predominance of the hydroxypyruvate decarboxylation (Figure 5). Furthermore, we did not see evidence for alternative explanations of this loss as metabolite concentrations of downstream products of glyoxylate NED (formate and folates) were similar between WT and *cat2*, suggesting that the glyoxylate decarboxylation reaction is unlikely to account for the excess CO_2_ release in *cat2* (Supplemental Figure 6). We also did not find evidence for elevated downstream products of serine decarboxylation as PC, PE and fatty acid profiles were similar between *cat2* and WT (Supplemental Figure 8).

There are other enzymatic decarboxylation reactions that have received attention recently that could help explain this increased CO_2_ loss. The import of glucose 6-phosphate (G6P) into the chloroplast could stimulate a G6P shunt that follows the oxidative branch of the pentose phosphate pathway around Calvin-Benson cycle and thus increasing CO_2_ release in the light (Sharkey and Weise 2016). It has been hypothesized that the G6P shunt could cause more CO_2_ release and lead to an increase in R_L_. Our data, showing no significant difference in R_L_ between WT and *cat2*, do not support that the excess CO_2_ release in *cat2* is due to additional CO2 release through the G6P shunt (Supplemental Figure 3). However, because R_L_ was determined under low CO_2_ concentrations and low light intensities, we could not exclude the possibility of excess CO_2_ released from G6P shunt under more physiological conditions. Alternatively, recent work highlights the potential for amino acid synthesis to contribute to CO_2_ release in a non-targeted metabolic analysis on sunflower showing that CO_2_ and O_2_ mole fraction changes the flux through several pathways involved in amino acid synthesis (Abadie and Tcherkez 2021). The CO_2_ release associated with these pathways was proposed to have potential impact on the amount of CO_2_ release per oxygenation, but it is difficult to evaluate this claim without quantitative flux estimates. Future labeling work combined with formal flux estimates should help resolve this source most conclusively (Xu et al. 2021), but is outside of the scope of the current work.

This work identifies a probable mechanism for excess CO_2_ release measured in mutants with perturbed photorespiratory metabolism but the presence of NED reactions during photorespiration may not be limited to mutant plants with a disrupted photorespiratory pathway (Cousins et al. 2008; Cousins et al. 2011; Keech et al. 2012). Photorespiration is impacted by changes in temperature in ways that could drive NED in WT plants. For example, the activity of glycolate oxidase increases more with temperature than catalase, possibly driving NED reactions (Grodzinski and Butt 1977). Additionally, post-translational modifications may modulate catalase activity, which could further modulate NED independently from protein content if activities were kept too low. For example, CAT2 contains a single phosphorylation site spanning residues 79-91 that is phosphorylated in response to nitrogen starvation (Hodges et al. 2013; Engelsberger and Schulze 2012). An increase in CO_2_ release per oxygenation under elevated temperatures would explain discrepancies in measurements of Γ* across many species, resulting in up to a 20% increase in the amount of carbon dioxide a plant loses per Rubisco oxygenation reaction (Walker and Cousins 2013). If present, this increase in CO_2_ released from photorespiration could present a potential route for improving the carbon recycling efficiency of photorespiration and subsequent net rates of CO_2_ fixation at elevated temperatures.

## Materials and methods

### Plant Material and Growth

*A. thaliana cat2* mutants (At4G35090, SALK 076998) were provided by Dr. Graham Noctor (Queval et al. 2007). WT and *cat2* plants used for measurements of Γ, Γ* and Φ_PS2_ were grown under a 12/12 day/night cycle at 90 μmol photons m^-2^ s^-1^ and 23/18 °C on a standard soil substrate A210 (Stender, Germany). WT and *cat2* plants used for PIB, membrane inlet and metabolic analysis were grown under a 11/13 day/night cycle at 100 μmol photons m^-2^ s^-1^ and 23/18 °C.

### Steady-state Gas exchange

Gas exchange was performed on the youngest, fully-expanded leaves of 4-6 week plants using a LI-6800 with a 3×3 cm measuring head (Li-COR Biosciences, Lincoln, Nebraska, USA). After measurements, leaf area enclosed by the cuvette was determined using the ImageJ FIJI distribution (Schneider et al. 2012; Schindelin et al. 2012). During all gas exchange measurements, leaf temperature was maintained at 25 °C and vapor pressure deficit was controlled at either 1 or 1.5 kPa for 25 °C. Measurements were performed in a climate-controlled chamber set to the measurement temperature.

Γ* values were measured using the common intersection method using slope-intercept regression (Laisk 1977; Walker and Ort 2015; Walker et al. 2016a) and light intensities of 250, 165, 120, 80 and 50 μmol photons m^-2^ s^-1^. No significant Kok effect was seen in light response curves measured at or above the light intensities used in Γ* measurements. Slope and intercept values were determined from the linear portion of CO_2_-response curves measured under each light intensity and CO_2_ concentrations between 10 and 3 Pa CO_2_ for determination of the common intersection value, which is equal to the intercellular CO_2_ concentration (C_i_^*^) at Γ*according to the equation Γ* = C_i_^*^ -R_L_ /g_m_ where R_L_ is determined from the y-axis value of the common intersection point and g_m_ was assumed to be 2.23 and 2.01 mol m^-2^ s^-1^ MPa^-1^ for 25 °C according to the temperature response measured previously in *A. thaliana* (Walker et al. 2013).

Light response curves for determining Φ_CO2_ values were measured under various CO_2_ and O_2_ environments controlled either using the native Li-COR 6800 functionality for CO_2_ or a synthetic N_2_ and O_2_ mixing system comprised of two mass flow controllers (red-y series, Vögtlin Instruments, Switzerland). Plants were acclimated under 250 μmol photons m^-2^ s^-1^ before being measured under 65, 50, 45, 40, 35, 30, 25 and 20 μmol photons m^-2^ s^-1^. For measurements of Φ_CO2_, the slope of CO_2_ assimilation vs absorbed light intensity was determined from Kok effect-free portions of the initial slope assuming a leaf absorbance of 0.843. For each condition, rates of *V*_*o*_ and the ratio *V*_*o*_*/V*_*c*_ were determined as described previously from the gas exchange measurements (Walker et al. 2014). Calculations of *V*_*o*_ and *V*_*o*_*/V*_*c*_ were made using chloroplastic CO_2_ concentrations calculated assuming a mesophyll conductance (g_m_) of 2.2 μmol m^-2^ s^-1^ Pa^-1^ from the intercellular CO_2_ concentrations measured at a point midway through the linear section of the light response curve (usually at 35 μmol photons m^-2^ s^-1^). While each light intensity across this range had slightly different *V*_*o*_ values, the relationship was linear overall, indicating that Φ_CO2_ appeared to be constant across the range and justifying a single representative *V*_*o*_ to be used (Fig. 3). Additionally, since the intercellular CO_2_ concentration was not greatly impacted across this range due to similar rates of net assimilation, *V*_*c*_/*V*_*o*_ was very constant across the light intensities used to determine Φ_CO2_. Respiration in the light was estimated according to the method of Kok (Kok 1948). Light-response curves were measured as sub-saturating light intensities. The rate of day respiratory (R_L_) was determined by the y-intercept by extending the part of light response curve after the compensation point to y-axis, removing any potential inflections due to the Kok effect.

### Membrane-inlet mass spectroscopy

Membrane-inlet mass spectroscopy was measured on leaf disks enclosed in a custom-built, thermostatted cuvette that allowed for inlet sampling and introduction of modified isotopic backgrounds (Cousins et al. 2008; Walker and Cousins 2013). The cuvette was built with multiple sampling and gas-release ports fitted with sampling septa. The membrane was composed of 0.005” fluorinated ethylene propylene film (CS Hyde Company, Lake Villa, IL, USA). The inlet line passed over a water trap maintained a few centimeters above liquid nitrogen. This was necessary to reach temperatures low enough to trap water at the low inlet pressures but not too low to freeze out carbon dioxide. The inlet line was passed into the ionizing source of a PrismaPlus Quadrapole Mass spectrometer (Pfeiffer Vacuum), which cycled through relevant masses for detection via an amplified faraday cup. The mass spec, roughing pump and turbo pump and time-resolved data collection software were provided by Bay Instruments (Port Easton MD, USA).

Measurements were made following a daily oxygen and carbon dioxide calibration and followed previous regimes and calculations (Cousins et al. 2008; Walker and Cousins 2013; Canvin et al. 1980). In brief, the leaf disk was placed within the chamber and sealed. The chamber was then flushed with nitrogen gas and ^18^O_2_ gas was injected to reach the desired atmosphere. Rates of dark CO_2_ release, ^18^O_2_ uptake and ^16^O_2_ release/leaks were monitored for ∼10 minutes before a custom-built LED light source was turned on. Rates of gas exchange were further monitored until a steady-state was reached and the chamber was injected with a saturating volume of ^13^CO_2_. Following this injection, ^12^CO_2_ release from photorespiration was monitored. A CO_2_ zero measurement was made before and after each experimental run by momentarily dipping the water trap into the liquid nitrogen. Time resolved mass spectrometer data was then processed by a pipeline of in-house python scripts to apply the necessary per volt calibrations and calculate the final rates of oxygen exchange, v_o_, v_c_ and ^12^CO_2_ release in a ^13^CO_2_ background. All processing scripts and chamber designs are available upon request.

### Post-illumination Burst (PIB)

The post-illumination burst of CO_2_ was measured using a LI-COR-6800 as described previously. During each measurement, the leaf was illuminated at 400 μmol photons m^-2^ s^-1^ for 40 min, and then darkened for 20 min. The total amount of CO_2_ release during the PIB was estimated as the area of PIB peak determined from the trace of CO_2_ release in the dark period after the baseline correction. The baseline was identified from the level of CO_2_ release in the last 200 s of dark period.

### Metabolic response to transient increases in rubisco oxygenation

Plants were treated with a rapid increase of light intensity from 50 μmol photons m^-2^ s^-1^ to 400 μmol photons m^-2^ s^-1^. Leaf tissues were collected before (time t=0) and after the shift to high light at the indicated time points. Samples were immediately frozen in liquid nitrogen and weighed before being stored at -80°C. Five biological replicates were performed for each genotype. The frozen samples were ground to a fine powder using a bead beating grinder with a sample holder containing dry ice. Metabolites were extracted with a solution of chloroform:methanol (3:7,v/v) and ribitol was used as internal standard. After centrifugation, the supernatant was freeze dried using a lyophilizer. Dried metabolites were methoximated (20 mg/mL methoxyamine in pyridine) and trimethylsilylated (MSTFA: TMSCI, 99:1) and then analyzed by GC-MS (Agilent 5975, GC/single quadrupole MS). GC-MS data were processed by Agilent MSD ChemStation. Metabolite derivatives were identified by comparison of the retention time with a known standard and comparison of the mass spectra with MS database. The amount of each metabolite was quantified by the total ion current signal of each metabolite peak normalized to the ribitol internal standard and tissue weight.

### Measurements of folates and formate

Formate was extracted by resuspending pulverized leaf tissue (∼100 mg) in 0.25 ml of 0.1 M HCl, with 10 μl of 10 mM amino butyric acid (ABA) added as an internal control. After centrifuging at 14,000 rpm for 20 min at 4 °C, the supernatant was collected, and the pellet was re-extracted with 0.25 ml of 0.1 M HCl. The supernatants were combined for analysis. Formate was analyzed using a published procedure (Xie et al. 2012) with some modifications. 50 μl of each sample was combined with 50 μl of Tetrabutylammonium bromide in Acetonitrile (20μmole/mL), 50 μl of Triethanolamine and 1 μl of 9-chloromethyl anthracene as a fluorescence-labeling reagent. The reaction was added up to 500 μl with acetonitrile and then was incubated at 75 °C for 50 min. After centrifuging at 14000 rpm for 10 min at 25 °C to precipitate the debris, the samples were separated on the Xterra MS C_18_ column (3.5μm, 4.6 x 100 mm, Waters, MA) with mobile phase of 64% acetonitrile and 36% water at the 1.0 mL/min constant rate.

Folates were extracted and analyzed as previously described (Hung et al. 2012). There were five biological replicates performed for each line for formate and folate analysis.

### Measurement of hydrogen peroxide

Hydrogen peroxide concentrations were measured in leaf tissues using a H_2_O_2_ Assay Kit from Abcam (ab102500, Cambridge, UK). Leaf tissues were harvested and homogenized as described above for metabolic response study, except that the frozen leaf tissues were immediately used for homogenization without storing at -80°C. Three biological replicates were performed for each genotype. After extraction and centrifugation, samples were deproteinized with 4 M perchloric acid, and then neutralized with 2 M KCl until pH between 6.5 and 8.0. All standards and deproteinized samples were incubated with OxiRed probe and horseradish peroxidase for 10 min at room temperature before measurements. Absorbance and fluorescence were measured with a 96-well plate reader (SpectraMax M2) at OD = 570 nm and Ex/Em = 535/587 nm.

### Lipid analysis

Polar glycerolipids were analyzed as described (Wang and Benning 2011). The Lipids were extracted from fresh leaves tissues. Polar lipids were separated on a silica-gel thin-layer chromatography plate treated with (NH_4_)_2_SO_4_ and a solvent system of acetone:toluene:water (91:30:7, v/v/v). Lipid spots were visualized with brief iodine vapor staining. Individual lipids, phosphatidylcholine (PC) and phosphatidylethanolamine (PE) were scraped off, and their fatty acid profiles were analyzed using gas-liquid Chromatography. Composition is presented as a mole percentage of total fatty methyl esters detected in each lipid. Three biological samples were collected for each line.

### Catalase activity

Catalase enzyme kinetics were determined on raw tissue extracts using an oxygen electrode by following the increase in oxygen production at various [H_2_O_2_] in a 50 mM potassium phosphate buffer, pH 8.1 to match the pH of a plant peroxisome (Rørth and Jensen, 1967; Switala and Loewen, 2002; Shen et al., 2013). The oxygen electrode temperature was set to 25 °C via a recirculating water bath.

### Determining rate constants and order of non-enzymatic reactions

The reaction between H_2_O_2_ and either glyoxylate or hydroxypyruvate was measured using UV-spectroscopy at 240 nm in a quartz reaction cuvette at 25 °C (Yokota et al., 1985) in a 50 mM potassium phosphate buffer, pH 8.1 to match the pH of a plant peroxisome. Since both glyoxylate and hydroxypyruvate also absorb at 240 nm, the absorbance drop attributed only to H_2_O_2_ decay was corrected by also accounting for the extinction coefficients of either glyoxylate (14 l mol^-1^ cm^-1^) or hydroxypyruvate (188.9 l mol^-1^ cm^-1^).

## Acknowledgements

We thank Marion Eisenhut for valuable discussions on photorespiration during the initial periods of planning this research. We additionally thank Ron Cook and Christoph Benning for help with the lipid analysis. We also thank Dr. Graham Noctor for providing the *cat2* seeds.

